# Rastair: an integrated variant and methylation caller

**DOI:** 10.64898/2026.03.19.712983

**Authors:** Zohar Etzioni, Liyuan Zhao, Pascal Hertleif, Benjamin Schuster-Boeckler

**Affiliations:** Ludwig Institute for Cancer Research, Nuffield Department of Medicine, University of Oxford, Oxford, UK; Softleif AB, Sweden

## Abstract

Cytosine methylation is a crucial epigenetic mark that impact tissue-specific chromatin conformation and gene expression. For many years, bisulfite sequencing (BS-seq), which converts all non-methylated cytosine (C) to thymine (T), remained the only approach to measure cytosine methylation at base resolution. Recently, however, several new methods that convert only methylated cytosines to thymine (mC→T) have become widely available. Here we present rastair, an integrated software toolkit for simultaneous SNP detection and methylation calling from mC→T sequencing data such as those created with Watchmaker’s TAPS+ and Illumina’s 5-Base chemistries. Rastair combines machine-learning-based variant detection with genotype-aware methylation estimation. Using NA12878 benchmark datasets, we show that rastair outperforms existing methylation-aware SNP callers and achieves F1 scores exceeding 0.99 for datasets above 30x depth, matching the accuracy of state-of-the-art tools run on whole-genome sequencing data. At the same time, rastair is significantly faster than other genetic variant callers, processing a 30x depth file takes less than 30 minutes given 32 CPU cores on an Intel Xeon, and half as long when a GPU is available. By integrating genotyping with methylation calling, rastair reports an additional 500,000 positions in NA12878 where a SNP turns a non-CpG reference position into a “de-novo” CpG. Vice-versa, rastair also identifies positions where a variant disrupts a CpG and corrects their reported methylation levels. Rastair produces standard-compliant outputs in vcf, bam and bed formats, facilitating integration into downstream analyses pipelines. Rastair is open-source and available via conda, Dockerhub, and as pre-compiled binaries from https://www.rastair.com.

## INTRODUCTION

Methylation of cytosine is a conserved epigenetic modification that plays crucial roles in regulating gene expression, cellular differentiation, and disease pathogenesis^1^. The ability to accurately measure the methylation state of cytosines across the genome is therefore essential for understanding biological processes and identifying disease biomarkers. In animals, methylation occurs predominantly at cytosines (C) followed by guanine (G), commonly referred to as CpG dinucleotides. The reverse-palindromic nature of CpG positions allows specialised methyl-transferases to identify hemi-methylated loci after DNA replication and re-establish full methylation, making it a heritable mark that is maintained across cell divisions. Notably, CpG dinucleotides are rare in mammalian genomes: in humans, they only constitute 2% of the genome, owing to the inherent mutability of mC as a result of replication errors and spontaneous deamination^2^.

For decades, bisulfite sequencing (BS-seq) has served as the gold standard for detecting methylated cytosine (mC) at single-base resolution. BS-seq works by chemically converting unmethylated C to uracil (U) which is then read as thymine (T), while leaving mC (and other cytosine modifications, such as hydroxymethyl-cytosine) intact^3^. After computational processing, C positions that do *not* show T can be assumed to have been methylated. However, BS-seq is a harsh chemical treatment that causes double-strand breaks and thus degrades a substantial portion of input DNA. This can result in low-complexity sequencing libraries and higher per-base error rates, especially when analysing samples with low amounts of input DNA.

Enzymatic methyl-seq (EM-seq) represents a more recent advancement that addresses some bisulfite sequencing limitations^4^. EM-seq uses a two-step enzymatic approach to detect mC positions: first, mC is oxidised to carboxyl-C (caC) by TET enzymes. In a subsequent reaction, APOBEC3A deaminates all C *except* caC to U. While EM-seq improves yield compared to BS-seq, both methods still suffer from issues relating to the loss of information as a result of converting nearly all genomic C to T: depletion of dTTPs and excess of dCTPs during sequencing by synthesis leads to low quality scores and higher error rates compared to conventional genome sequencing^3^.

Subsequent alignment to the reference genome requires specialised tools that take significantly longer to run, and a higher fraction of reads cannot be accurately mapped, owing to the loss of distinguishing features after conversion of most C to T.

These limitations have prompted the development of alternative methodologies that selectively convert *only* mC to T, which we will refer to as “mC→T” methods below. We were involved in the development of TAPS, a method that leverages TET enzymes to oxidise mC to caC, followed by a chemical conversion of caC to di-hydroxy-uracil (DHU), which is then read as T^5^. Later, Wang *et al*. described DM-seq, an approach that uses two enzymes to selectively convert only mC to T^6^. More recently, Illumina also released a fully enzymatic method they call 5-Base sequencing (5-Base) which leverages a proprietary enzyme to directly convert mC to T^7^. As mC→T methods result in reads that contain few chemically-induced mismatches against the reference genome, they can be conveniently aligned with non-specialised high-performance aligners like bwa-mem2^8^. The higher information content per read reduces the number of unaligned or mis-aligned reads, resulting in higher total mapping rates and more even coverage across the genome compared to BS-seq and EM-seq^5^.

In addition to being the site of methylation, cytosines in CpG positions are also frequently mutated to thymine^2^. Consequently, CpGs accrue more single-nucleotide polymorphisms (SNPs) than any other sequence context^9^. In fact, it has been shown that a large number of common SNPs can both create or interrupt a CpG dinucleotide, which can have significant cis-regulatory effects on neighbouring CpGs^10,11^. As C>T SNPs can be mistaken for chemical conversion in all mC→T methods, as well as in BS-seq and EM-seq, it is therefore important to not only quantify the methylation state of cytosines, but also to detect genetic variants in the same sample. Several algorithms have been devised to call genetic variants from BS-seq and EM-seq data^12,13^. However, their sensitivity and specificity lags behind that of common variant callers on WGS data^14,15^. Consequently, few studies incorporate genotype information from epigenetic sequencing data into their analyses.

Here, we present *rastair*, an integrated toolkit that combines genetic variant detection with methylation quantification from mC→T data. We present evidence that rastair outperforms existing variant callers for BS-seq and mC→T methods in both variant calling accuracy and computational performance. Unlike most other methylation callers, rastair adjusts the estimated methylation depending on patient genotype for C>T SNPs at CpG sites and clearly reports “de-novo” CpG positions that are formed by SNPs, thus reporting nearly 500,000 additional CpG positions - relative to the reference sequence - for a typical human genome.

## METHODS

### SOURCE DATA

#### TAPS+

Genomic DNA was derived from human reference cell-line NA12878/HG001^16^. Libraries were generated by Watchmaker Genomics using the TAPS+ kit and sequenced on an Illumina NovaSeq X Plus. Processing of reads was performed by us: reads were trimmed with trimGalore 0.6.10 and aligned using bwa-mem2 2.3.0 to the hg38 reference genome. After alignment, duplicates were marked with Picard MarkDuplicates (v4.5.0.0).

#### 5-BASE

Illumina 5-Base sequencing data of NA12878 were downloaded from Illumina Basespace, including aligned bam files, vcf.gz files and CX files. According to the file headers, these were generated with the DRAGEN methylation pipeline version 4.4.1.

### SNP EVALUATION

#### REFERENCE DATA

The GRCh38 human reference genome was obtained from the Broad Institute GATK resource bundle (gs://genomics-public-data/resources/broad/hg38/v0) and supplemented with HPV, HBV, and HCV spike-in sequences. High-confidence SNP calls and confident genomic regions for HG001 (NA12878) were obtained from the Genome in a Bottle Consortium (GIAB) v4.2.1 release (ftp://ftp-trace.ncbi.nlm.nih.gov/ReferenceSamples/giab/release/NA12878_HG001/latest/), comprising the benchmark VCF and high-confidence region BED files for GRCh38 chromosomes 1–22. Genome stratification regions (v3.3) were obtained from the GIAB stratification resource (ftp://ftp-trace.ncbi.nlm.nih.gov/ReferenceSamples/giab/release/genome-stratifications/v3.3/), enabling separate assessment of caller performance in genomic contexts including tandem repeats, low-mappability regions, segmental duplications, and extreme GC content. GATK resource bundle files used for base quality score recalibration included dbSNP build 138, Mills and 1000 Genomes gold-standard indels, and known indels for GRCh38 (https://console.cloud.google.com/storage/browser/genomics-public-data/resources/broad/hg38/v0).

#### INPUT ALIGNMENTS

Standard whole-genome sequencing data for HG001 (NA12878) at approximately 300x coverage were obtained from the GIAB consortium (ftp://ftp-trace.ncbi.nlm.nih.gov/ReferenceSamples/giab/data/NA12878/NIST_NA12878_HG001_HiSeq_300x/). This dataset was generated on the Illumina HiSeq 2500 in Rapid Mode (v1) using 2×148 bp paired-end reads with TruSeq DNA PCR-Free library preparation and approx. 550 bp insert size (SRA accessions SRX1049768–SRX1049855).

#### DOWNSAMPLING

To evaluate SNP calling performance as a function of sequencing depth, the high coverage alignments were systematically downsampled to target coverages of 45x, 30x, 10x, and 5x using `samtools view -bs` (samtools v1.21). Mean coverage was computed exclusively within GIAB high confidence regions using samtools depth -b, ensuring that downsampling fractions accurately targeted the benchmark-relevant portion of the genome. At each target coverage, three independent replicate BAMs were generated using random seeds 42, 43, and 44, to quantify stochastic sampling variability. Read group tags were added to each BAM using `samtools addreplacerg` to satisfy downstream caller requirements. In total, 12 WGS and 12 TAPS+ downsampled BAMs were produced.

#### WGS VARIANT CALLING

Four variant callers were applied to the WGS data, all allocated 32 CPU threads:

**GATK HaplotypeCaller** (v4.6.2.0): Base quality score recalibration (BQSR) was performed using `BaseRecalibrator` and `ApplyBQSR` with dbSNP 138, Mills and 1000 Genomes gold-standard indels, and known indels as known-sites resources. Variants were called with `HaplotypeCaller` using `--native-pair-hmm-threads 32`. SNPs were selected using `SelectVariants -select-type SNP` and hard-filtered following GATK best practices: `QD < 2.0, QUAL < 30.0, SOR > 3.0, FS > 60.0, MQ < 40.0, MQRankSum < -12.5, ReadPosRankSum < -8.0`.

**DeepVariant** (v1.9.0): Run via Singularity container with the standard WGS model `(--model_type=WGS)`.

**Octopus** (Invitae fork, build eae1ab48): Run via Singularity container on chromosomes 1–22, X, Y, and M with `--ignore-unmapped-contigs` and `--threads 32`.

**Clair3** (Singularity container): Run with the Illumina platform model (`--platform=“ilmn”, --model_path=“/opt/models/ilmn”`).

#### TAPS VARIANT CALLING

3 methylation-aware variant callers were applied to TAPS+ data, each allocated 32 CPU threads:

**Rastair** (v2.1.0-rc1 compiled from Rust source in release mode): Called with `rastair call --threads 32 --vcf-info-fields AS_SB`. Only PASS-flagged SNPs were considered.

**TVC** (TAPS+ Variant Caller; Singularity container): A TAPS-aware variant caller run with minimum base quality 20, minimum mapping quality 1, and minimum depth 2. Output was filtered to SNPs only; no GQ or PASS filter was applied.

**Biscuit** (Singularity container): A bisulfite-sequencing variant caller (`biscuit pileup`) applied experimentally to TAPS+ data, noting that Biscuit is designed for bisulfite (unmethylated C→T) rather than TAPS+ (methylated C→T) conversion chemistry.

#### VCF NORMALISATION

All query and truth VCFs were processed through an identical normalisation pipeline using bcftools v1.21: multiallelic records were decomposed into biallelic sites (`bcftools norm -m-any`), left-aligned against the reference, and reference allele mismatches flagged with warnings (`--check-ref w`, expected due to viral spike-in sequences in the reference). The output was restricted to SNP type variants (`bcftools view -v snps`) with PASS filter status where applicable.

#### BENCHMARKING

SNP calls were benchmarked against the GIAB v4.2.1 high-confidence truth set using hap.py v0.3.12 (Illumina) with the vcfeval engine (RTG Tools), which performs haplotype-aware variant comparison without requiring phasing information. Evaluation was restricted to PASS-flagged variants (--pass-only) within GIAB high-confidence regions on chromosomes 1–22, X, and Y. Stratified analyses were performed using the GIAB v3.3 genome stratification regions, with key stratifications including non-difficult regions (high mappability, no repeats), all difficult regions (union of low-mappability, segmental duplications, and tandem repeats), low-mappability regions, and tandem repeat regions.

Precision 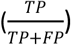, recall 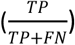and F1 score (harmonic mean of precision and recall) were computed from hap.py output. Reproducibility across the three replicate downsamples was quantified using the coefficient of variation (CV) of F1 scores. False positive rate 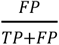 and false negative rate 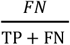 were computed separately to characterise caller error profiles.

### METHYLATION EVALUATION

#### METHYLATION CALLERS

The aligned BAM file for a 5-Base NA12878 sample was downloaded from Illumina BaseSpace. Three methylation callers were applied to the same BAM file to enable methylation evaluation:

**Rastair**: Rastair v2.0.0 was run with default parameters. Methylation information for both reference and variant cytosines was reported in the BED and VCF outputs, which were internally consistent.

**DRAGEN** (v4.4.1): DRAGEN output files were obtained from Illumina Basespace. These included a cytosine methylation report (CX_report.txt.gz) containing per reference cytosine methylation status, and a VCF methylation report (vcf.gz) containing variant-associated methylation estimates. Details about DRAGEN 5-Base outputs can be found at https://help.dragen.illumina.com/product-guide/dragen-v4.4/dragen-methylation-pipeline/dragen-5base-pipeline

**MethylDackel** (v0.6.1): Run with default parameters. The output cytosine report (cytosine_report.txt.gz) was used for downstream analysis.

#### OUTPUT HARMONISATION AND CONTEXT FILTERING

Evaluation was restricted to chromosomes 1-22, X, and Y. For rastair, CpG calls were extracted from the BED file. For DRAGEN and MethylDackel, only cytosines in CpG (CG) context were retained from the cytosine report files unless otherwise noted. Since MethylDackel reports all reference CpG positions regardless of coverage, only sites with coverage were included in the analysis. Filtered CpG positions were then categorised as either shared between tools or unique to a specific tool.

#### REFERENCE METHYLATION CALLING

Reference CpG regions were defined using rastair as sites labelled `REF` with genotype C/C or G/G, indicating no variant affecting the CpG context. For rastair, methylation levels were retrieved from the `beta_est` field in the BED file. For DRAGEN and MethylDackel, methylation levels were calculated as:

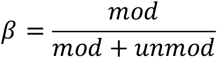

where *mod* and *unmod* represent methylated and unmethylated read counts reported in the cytosine report files. Only reference CpG positions identified by all three callers were included in the concordance analysis of reference methylation.

#### VARIANT METHYLATION CALLING

SNPs were extracted from the rastair and DRAGEN VCF files using `bcftools view -v snps`. For DRAGEN, variant-associated CpG methylation estimates were obtained from the VCF file in accordance with the DRAGEN 5-Base specification. Specifically, methylation levels were extracted from the `FORMAT:M5mC` field based on the context annotations in `INFO:M5mC`. Only entries marked with `z` were considered. Methylation values were matched positionally to their corresponding context codes. When multiple `z` annotations were present at a given locus, the first corresponding methylation value was selected as the representative estimate for downstream analysis. De-novo CpGs identified by rastair were further classified into de-novo with variant, defined as positions lacking an alternative allele entry in the ALT field, and de-novo without variant.

### OESOPHAGEAL CANCER SAMPLE PROCESSING

#### SAMPLE COLLECTION AND SEQUENCING

Samples were collected as part of the LUD2015-005 clinical trial as described in Carroll *et al*.^17^. Briefly, genomic DNA was extracted from fresh-frozen endoscopic biopsies. Patient-matched samples of tumour-free duodenum were used as control. DNA extracted from these fresh biopsies was split, with one part being processed according to the protocol detailed in Liu et al ^5^, the other undergoing standard WGS library preparation as previously described^17^. Sequencing was performed on Illumina NovaSeq 6000 and NovaSeq X platform.

#### DATA PROCESSING

TAPS reads were trimmed, aligned and duplicates marked as described above. WGS data from control samples was processed as previously described^17^. We used Octopus v0.7.0 to call germline variants using default settings. Rastair v2.0.0 was used to call methylation and variants from tumour TAPS samples. Figures were generated in R using custom scripts.

## RESULTS

### IMPLEMENTATION OF AN INTEGRATED METHYLATION AND VARIANT CALLER

#### SNP CLASSIFICATION AND FILTERING

Rastair processes genomic loci sequentially and independently, which allows for efficient parallelisation (Figure 1A, Supp. Figure 1A&B). Four types of loci are identified and further processed:

1. CpG sites in the reference genome
2. CpG sites in the reference that show evidence of a variant that cannot be explained by chemical conversion *(“CpG” variants)*
3. Evidence of a variant at non-CpG positions which would result in a CpG position, e.g. a T>C in a TpG dinucleotide *(“de-novo CpG” variants)*
4. Any other non-CpG variant *(“other” variant)*

**Figure 1.**
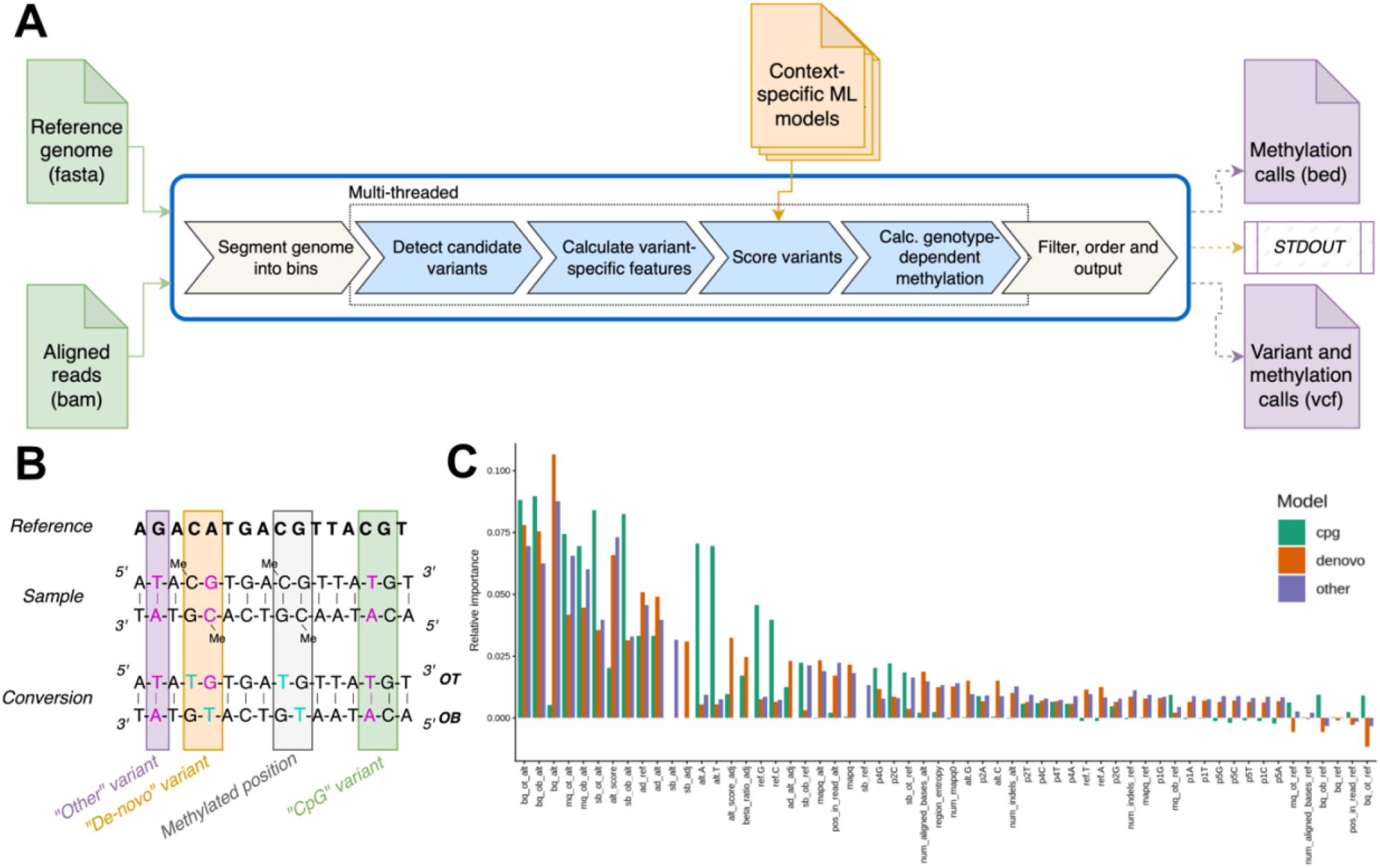
**(A)** Schematic outline of the rastair variant and methylation calling process flow. **(B)** Schematic of different variant classes observed in mC→T data: In addition to “regular” variants (here called “other”), variants can create a new CpG position relative to the reference (“de-novo variant”) or delete a CpG position (“CpG variant”). After conversion, each of them shows distinct sequence patterns in original top (OT) and original bottom (OB) reads. **(C)** Feature importance for the three different models used to classify variants. Importance was estimated by permuting each feature and measuring drop in prediction accuracy. “sb”: strand bias, “adj”: adjacent position, ie the G or C to a C or G, respectively, in a CpG or de-novo CpG. “bq”: RMS base quality. “mq” RMS mapping quality

For CpGs that show no variant evidence, the pipeline simply reports methylation by counting the evidence for C and T on the relevant strand (Figure 1B) and calculating β as 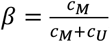 where *c*_*M*_ and *c*_U_ denote the number of reads (passing any pre-processing filters, such as minimum mapping quality) that show evidence for methylation and no methylation, respectively.

When variant evidence is present at a position, the workflow gathers a comprehensive set of contextual features that are then fed to one of three machine learning models to classify the observed variant as real or artefactual. Across all models, the features share several common components:

- the reference base and the alternate base, each represented with one-hot encoding (four features per base)
- the sequence context, comprising the reference bases two positions before and after the current position, also one-hot encoded
- the entropy of the surrounding 100-base region around the position
- the root-mean-square (RMS) of base qualities for the reference and alternate allele, both overall and per strand
- the RMS of mapping qualities of reads supporting the reference and the alternate allele, both overall and per strand
- the fraction of reads with mapping quality zero, relative to all reads at that locus
- the fraction of reads that show evidence for the ref and the alt alleles, relative to the total number of reads at this locus, both overall and split by strand
- the RMS of the number of insertions or deletions per read for the reference and alternate allele
- the RMS of the distance of the variant from the nearest end of all reads supporting the reference and the alternate allele

In addition to these common features, we also capture features unique to potentially methylated loci. At CpG position, we calculate a base-quality weighted score for the enrichment of alternate over reference reads:

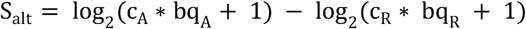

where c_A_ and c_R_ denote the counts of reads on the non-modified strand with evidence for the reference and the alternate allele, respectively, and bq_A_ and bq_R_ denote the respective base quality scores. The CpG and de-novo CpG models also include information about the adjacent G or C position, respectively. This acknowledges that methylation is commonly symmetrical, so evidence for or against chemical conversion at a neighbouring position influences the likelihood of an adjacent variant call. In particular, we the log ratio of methylation beta values between the current position and an adjacent CpG position and the S_A_ of the adjacent position.

### GENOTYPE CLASSIFICATION

These features were used to train a random forest model. After hyperparameter tuning, we settled on 800 trees and two features per split. As training data, we used 8000 true and 20000 false variants each per model, randomly sampled out of all candidate variants of each type identified on chromosome 12 of TAPS+ sequenced DNA from NA12878 (see Methods). The train/test split was chosen to better reflect the class imbalance between true SNP and true non-SNP positions. We then used Platt scaling to calibrate each model against the remaining variants on chromosome 12 of the respective type. This calibration results in scores that can be interpreted probabilistically: values above 0.5 represent a larger than 50% likelihood of being a true positive.

For each variant position whose ML score exceeds the chosen threshold for inclusion (0.5 by default), we subsequently estimate the genotype zygosity using a straightforward heuristic. We reasoned that any real variant at a diploid position is either homozygous or heterozygous. To distinguish the two, we calculate two probabilities given the observed read count data *D*:

1. *P*(*D* | *hom*) that the variant is homozygous, and any evidence for heterozygosity is due to sequencing error, *i.e*. the number of observed reference and alternate alleles follows a binomial distribution with *p* = *ϵ, q* = 1 − *ϵ*, where *ϵ* is the estimated sequencing-platform-specific base calling error rate.
2. *P*(*D* | *het*) that a variant is heterozygous, *i.e*. the number of observed reference and alternate alleles follows a binomial distribution with *p* = 0.5, *q* = 0.5.

The classification is then made as *M* = *argmax*_*M*∈{*hom,het*}_*P*(*D* | *M*). At CpG and de-novo CpG positions, we count potentially converted reads as evidence for the C or G allele, respectively, unless this is ambiguous with the alternative allele, in which case we ignore all reads on the strand that would be affected by chemical conversion. For example, imagine a C to A SNP at a CpG dinucleotide: here, T on the OT strand is most likely evidence for a C allele. In contrast, at a C to T SNP, we cannot distinguish T due to conversion from genetic T on the OT, so we ignore all OT reads for the purpose of genotype classification.

### GENOTYPE ADJUSTED METHYLATION ESTIMATES

We always report both the C and G position for any methylated position, including for de-novo CpGs that pass the ML threshold. However, the presence of a methylatable CpG can be allele dependent. Rastair does not yet perform phased methylation calling. Instead, we currently adjust the estimated beta value of CpG loci affected by a SNP.

For homozygous SNPs that would disrupt a CpG, we always return *β* = 0 at the variant position.

For heterozygous SNPs at C or G positions, only half of the reads are expected to be derived from the methylatable allele. This is a particular issue at C to T (and G to A) SNPs, where the “genetic” T/A could be mis-counted as chemical conversion. We therefore adjust the estimated methylation at these positions using the following equation:

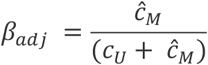

where

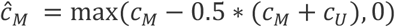

Here, *ĉ*_*M*_ represents the “excess” of methylated reads above what is expected to be reads from the non-methylated allele.

### VARIANT CALLING ACCURACY

We benchmarked rastair’s accuracy using the Genome-In-A-Bottle reference variant set for the human cell line NA12878. We sequenced DNA from NA12878 cells to 50x with and without TAPS+ conversion (see Methods). We then sub-sampled the data to different depth (from 5x to 45x) and called SNPs. On non-TAPS WGS data we tested four widely used algorithms: DeepVariant^18^, Octopus^19^, GATK HaplotypeCaller^20^ (with hard cutoffs), and Clair3^21^ (see Methods). For TAPS+ data, we tested TVC (developed by Watchmaker Genomics) and Biscuit, a tool commonly recommended for variant calling on whole-genome bisulfite sequencing (WGBS) data.

Rastair consistently outperforms other methylation-aware variant callers (TVC, Biscuit) and achieves variant calling accuracy close to that of existing variant callers on WGS data, especially at sequencing depth above 10x (Figure 2A). Specificity of TAPS-based callers was observed to be more coverage dependent than for WGS-based callers (Figure 2B, left). In contrast, rastair exhibited higher sensitivity/lower false-negative rates at 5x and 10x coverage than e.g. Clair3 and DeepVariant (Figure 2B, right). We also compared variant calling accuracy of rastair on Illumina 5-Base data sequenced to 45x depth and compared it to variant calls generated by DRAGEN (see Methods). In the high-confidence regions used in Fig 2, rastair achieved an F1 score of 98.9%, compared to 99.4% for DRAGEN. Across all available reference variant calls for NA12878 (ie including lower-confidence calls), rastair had an F1 of 90.6% compared to 89.9% for DRAGEN (Supp. Fig 2).

**Figure 2.**
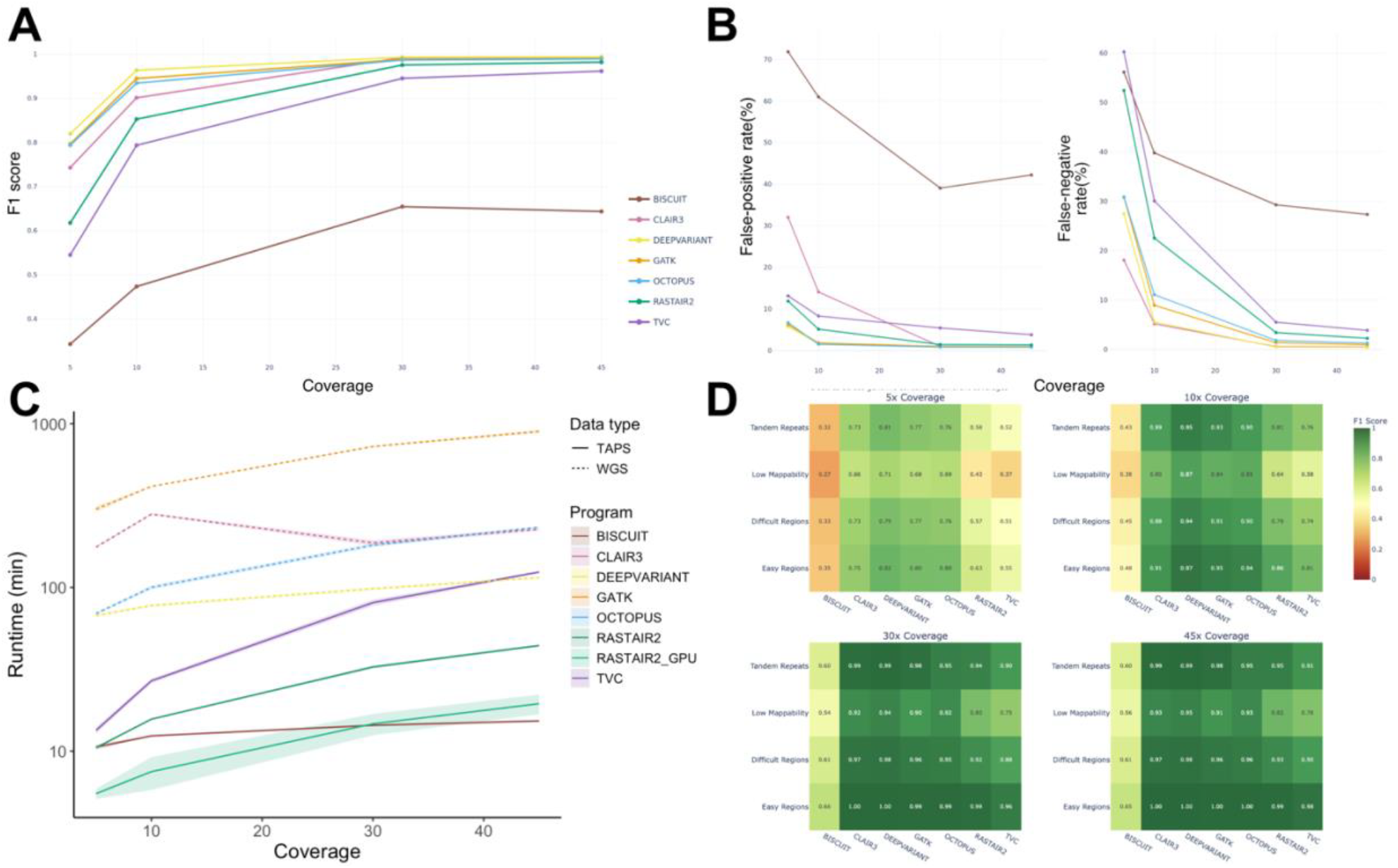
**(A)** F1 score at increasing coverage of different variant callers on WGS (Clair3, DeepVariant, Octopus, GATK HaplotypeCaller) or TAPS+ input data (rastair, TVC, Biscuit), calculated using hap.py, restricting to high-confidence regions of the GIAB reference call set. **(B)** False positive (left) and false negative (right) rates at increasing coverage by caller. **(C)** Absolute runtime of callers with input of increasing coverage depth. Each caller was run 3 times and given 32 CPU cores on Intel Xeon Gold CPU with 2.6GHz. Standard deviation is denoted as transparent ribbon. **(D)** F1 score of callers stratified by coverage and type of genomic region: Tandem Repeats, regions of low mappability, and regions classified as “easy” or “difficult” by the GIAB consortium.

We performed repeat runs of each caller on a dedicated compute node with 2.6GHz Intel Xeon Gold CPUs with 32 cores to measure the time required to call variants on a full genome of the given sequencing depth. Rastair is faster than all evaluated callers except Biscuit (Figure 2C). To further improve runtime, we also implemented GPU-acceleration of the ML-based variant classification step of rastair (see Supplementary Methods). Using GPU acceleration approximately halves the runtime compared to the CPU version of rastair: processing a 45x coverage bam file with rastair took 45 minutes with the CPU version using 32 cores compared to only 19 minutes when using GPU acceleration (Supp. Table 1). In contrast, TVC took nearly two hours (118 minutes), and Octopus required close to 4 hours (238 minutes) to process the same file.

### COMPARISON WITH OTHER METHYLATION-CALLERS

While the “Genome In A Bottle” (GIAB) consortium provides deeply characterised reference variants for the NA12878 cell line, there is - to our knowledge - no reference dataset with definitively validated methylation levels at specific CpG positions. In fact, given the dynamic nature of DNA methylation, it would be difficult to establish one. In order to evaluate the performance of rastair, we therefore resorted to comparing our methylation calls to those produced by two other tools that report methylation levels while accounting for potential effects of genetic variation: MethylDackel and the newly released DRAGEN methylation pipeline.

Aligned BAM files for one exemplary 5-Base dataset, together with matching DRAGEN CX report and VCF files, were downloaded from Illumina BaseSpace. MethylDackel and rastair were run with default parameters. MethylDackel and DRAGEN CX report files were initially filtered for “CG” context only. Rastair reports nearly 1.9 million more CpG positions than MethylDackel or DRAGEN’s CX report file (Figure 3A). Positions shared between MethylDackel and rastair consist predominantly of homozygous variants affecting reference CpG positions.

**Figure 3.**
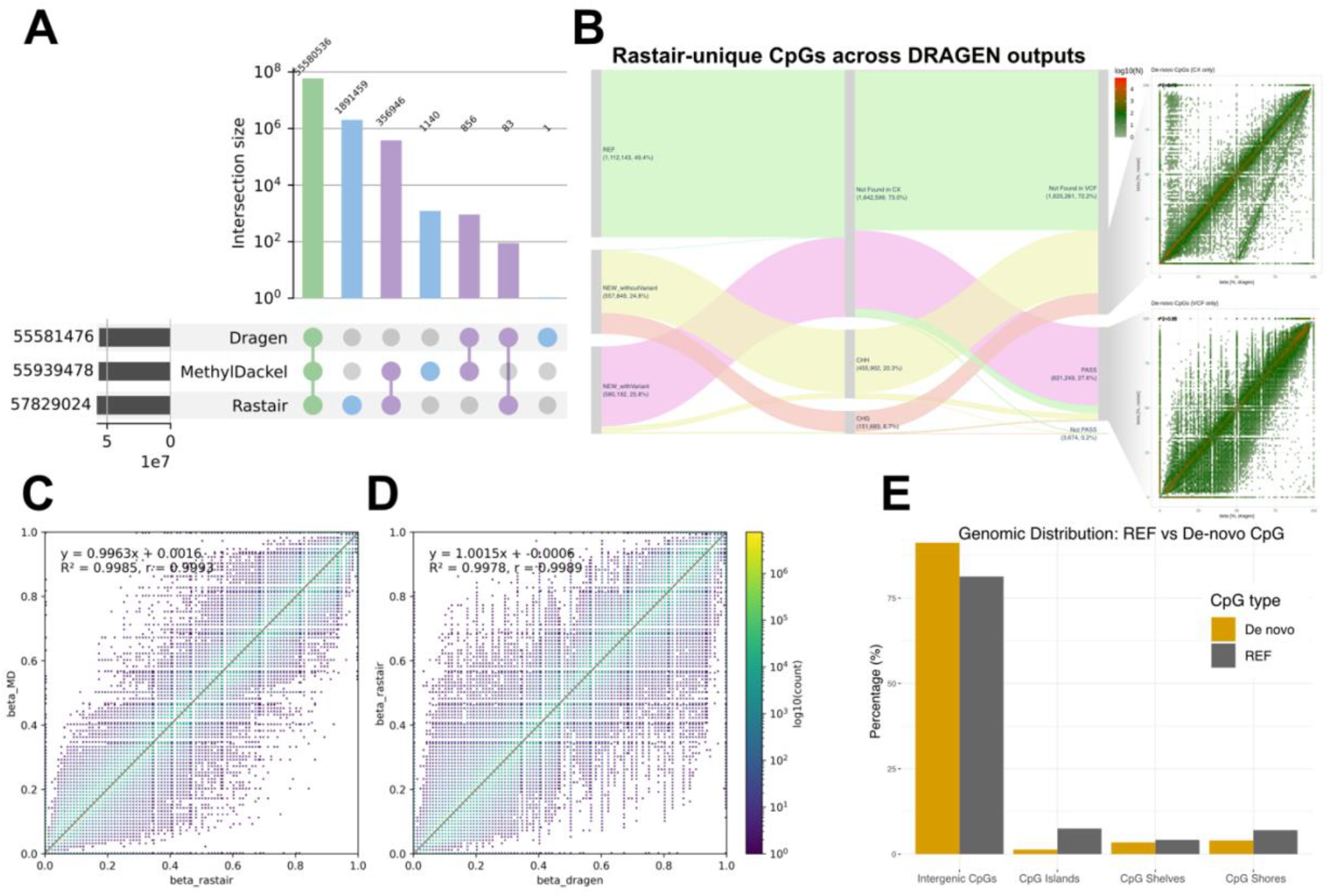
**(A)** Upset plot showing combinations of CpG positions with methylation calls from rastair, DRAGEN and MethylDackel. The horizontal bars show the total number of called CpG positions per method. **(B)** Sankey plot showing which CpG positions unique to rastair are reported as CpH methylation in the DRAGEN’s CX report and/or as CpG methylation in the DRAGEN vcf file. **(C, D)**: pairwise comparison of estimated β values for overlapping CpG positions, excluding de-novo CpG positions and positions affected by SNPs, for rastair vs MethylDackel (C) and rastair vs DRAGEN (D). **(E)** Distribution of de-novo CpGs across genomic features, compared to reference CpGs.

MethylDackel performs no beta correction on these, while DRAGEN seems to entirely remove them from the report. In contrast, rastair assigns them a β value of 0 to reflect that these sites are covered but no methylation is present.

Focussing on reference CpG positions that are unaffected by genetic variants, we observe high concordance between all tools, with *R*^*2*^ values above 99% for both MethylDackel and DRAGEN (Figure 3C,D). The observed variation is a result of differences in the default filtering of reads and bases: for example, MethylDackel sets different thresholds for base- and mapping-quality compared to rastair (rastair: min mapQ 1 vs 10 in MethylDackel, rastair min baseQ 10 vs 5 in MethylDackel). When we ran rastair with the same filter settings, the R^2^ increases to 99.98% (Supp. Figure 3). The exact filter settings for DRAGEN are unknown, but it is notable that rastair and MethylDackel share higher concordance while DRAGEN estimates higher beta values than rastair in a subset of positions (Figure 3D).

Amongst the sites that are unique to rastair, half are reference CpG positions that were not reported by either MethylDackel or DRAGEN due to mapping quality, base-quality and/or coverage thresholds (Fig 2B). The other half are de-novo CpGs that are formed when a SNP creates a new CpG allele. In total, we detect over 500k such new CpG dinucleotides relative to the reference. These de-novo CpGs can be found in CpG islands, shelves and shores, however they are enriched in intergenic CpGs and depleted in CpG islands (CGIs) (Figure 3E), most likely as a result of the lower mutation rate of un-methylated CpGs in CGIs^22^.

For de-novo CpGs, rastair reports both the C and the G component in both the bed and vcf output files, as methylation is usually symmetrical and information from the adjacent base can be valuable in interpreting the methylation state of a locus. In contrast, de-novo C or G positions are not reported in the DRAGEN CX file but only in the DRAGEN vcf output (Figure 3B, red stream). Vice-versa, the adjacent non-variant base is not included in DRAGEN’s VCF file, but can often be found as non-CpG methylation in the CX file (Figure 3B, yellow stream).

DRAGEN thus splits methylation information for the same locus across two files. Furthermore, the β estimates provided by DRAGEN in their vcf file are corrected for the zygosity of the underlying variant and thus correlate closely with rastair’s estimates (Figure 3B, bottom right). Conversely, the estimates in the CX file are uncorrected and exhibit a strong skew where genetic T/A at heterozygous C>T and G>A variants is counted as methylation, leading to artificially high β values (Figure 3B, top right).

### DETECTING GENOTYPE-METHYLATION RELATIONSHIPS WITH RASTAIR

Rastair’s ability to detect genotype at the same time as methylation can be leveraged to directly detect effects of genotype on gene regulation (Fig 4A). A good example of such a link is rs1800734, a rare G>A variant in the promoter of MLH1, a core gene in the mismatch-repair pathway. Hypermethylation of a CpG island in the MLH1 promoter is associated with microsatellite instability (MSI) and increased cancer risk^23^. It has previously been reported that rs1800734 is linked to hypermethylation of the MLH1 promoter and thus increases the risk of MSI^24^. We hypothesised that rastair would be able to identify this type of genotype-epigenome relationship from TAPS sequencing data. To test this, we leveraged a dataset of 24 tumours from oesophageal cancer patients that were sequenced with TAPS to an average of 48x coverage. We also performed somatic variant calling on these tumours from paired tumour and normal tissue samples.

**Figure 4.**
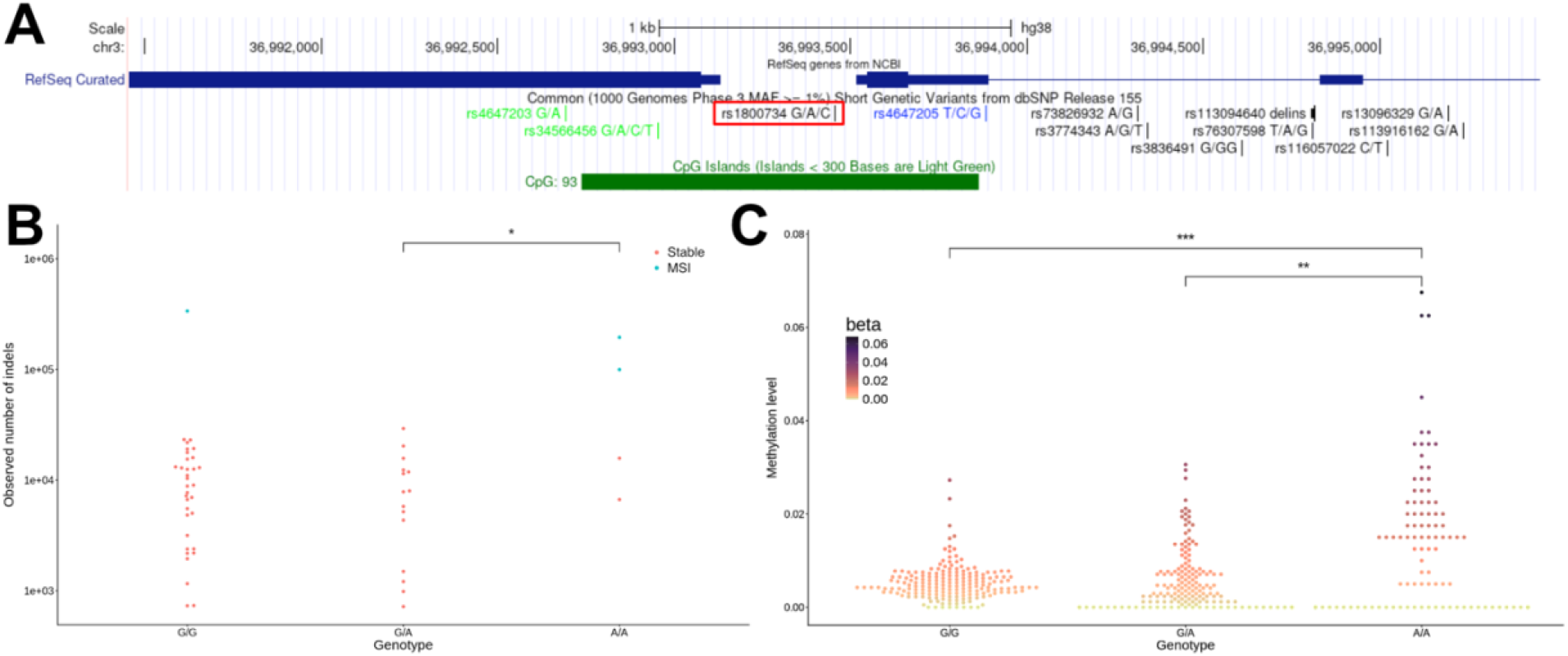
**(A)** Overview of the MLH1 promoter region. A 1.1kb CpG-island covers parts of exon 1 of EPM2AIP1 (left) and the exon 1 of MLH1 (right). The position of rs1800734 is highlighted in red. **(B)** Number of somatic insertion/deletion mutations observed in tumour samples, stratified by rs1800734 genotype. Samples classified as “MSI” are highlighted in blue. Significance by Wilcoxon pairwise ranksum test. **(C)** β values at CpGs in the MLH1 CpG-Island per patient, stratified by rs1800734 genotype. Significance by Wilcoxon pairwise ranksum test.

Based on somatic variant calls, we observed MSI in 3 patients (see Methods). Using rastair, we genotyped the rs1800734 locus in all samples. All rastair calls matched the genotype called by Octopus on adjacent normal tissue WGS. 4 sample were homozygous for rs1800734 (*ie*. carried two A alleles), two of which were called as MSI (p=0.013, Fisher’s Exact Test; Figure 4B). Methylation levels at the MLH1 promoter were significantly elevated in patients with a homozygous A/A genotype (Figure 4C), supporting the notion of a direct link between rs1800734, epigenetic silencing of MLH1 and MSI.

### OTHER OUTPUTS

In addition to per-position methylation and genotype information, rastair also provides per-read output in multiple different formats. Our own custom text-based format reports, for each sequenced read, its genomic location, the total number of CpGs covered by the read, how many of them were observed as methylated, and the position in the read that represent the modified and unmodified bases. This format is convenient to e.g. calculate *α*-values (*ie*. the fraction methylated bases per read), and for other applications that require read-level methylation information. Furthermore, rastair can also annotate the original bam file with per-read methylation tags. We support both the SAM-standard way of reporting methylation via the `MM` tag, as well as Bismark’s `XM/XR/XG` tags.

Rastair also provides a companion script to generate a number of QC figures for mC→T data as an easy to interpret HTML report. This comprises an M-bias plot as first described by Hansen *et al*.^25^ (Figure 5A). This plot is useful to identify technical issues at the ends of reads, *eg*. due to incomplete adapter trimming or loss of methylation due to end-repair. Rastair also suggests soft-clipping parameters that can be used with the `--nOT` and `--nOB` parameters of `rastair call` to exclude read ends from methylation calling but retain them for variant detection. However, we noticed that M-bias plots are sometimes insufficient to capture the technical issues present in certain data types. For example, cell-free DNA is naturally fragmented into short pieces of DNA with frequent “jagged ends” ^26^. This can lead to a significant drop in methylation when the single-stranded DNA is re-synthesized during end-repair. Due to the unique features of cfDNA, this leads to a fragment-length-dependent pattern of conversion that can be captured in a “V-bias” plot (Figure 5B). Here, the average methylation of CpGs at the respective position in the original DNA fragment is represented by colour, and the fragments are ordered by their original length (as estimated by the insert size after alignment). This illustrates that in cfDNA, the latter part of long fragments is frequently re-synthesized during library preparation, reducing the informativeness of the second read pair.

**Figure 5.**
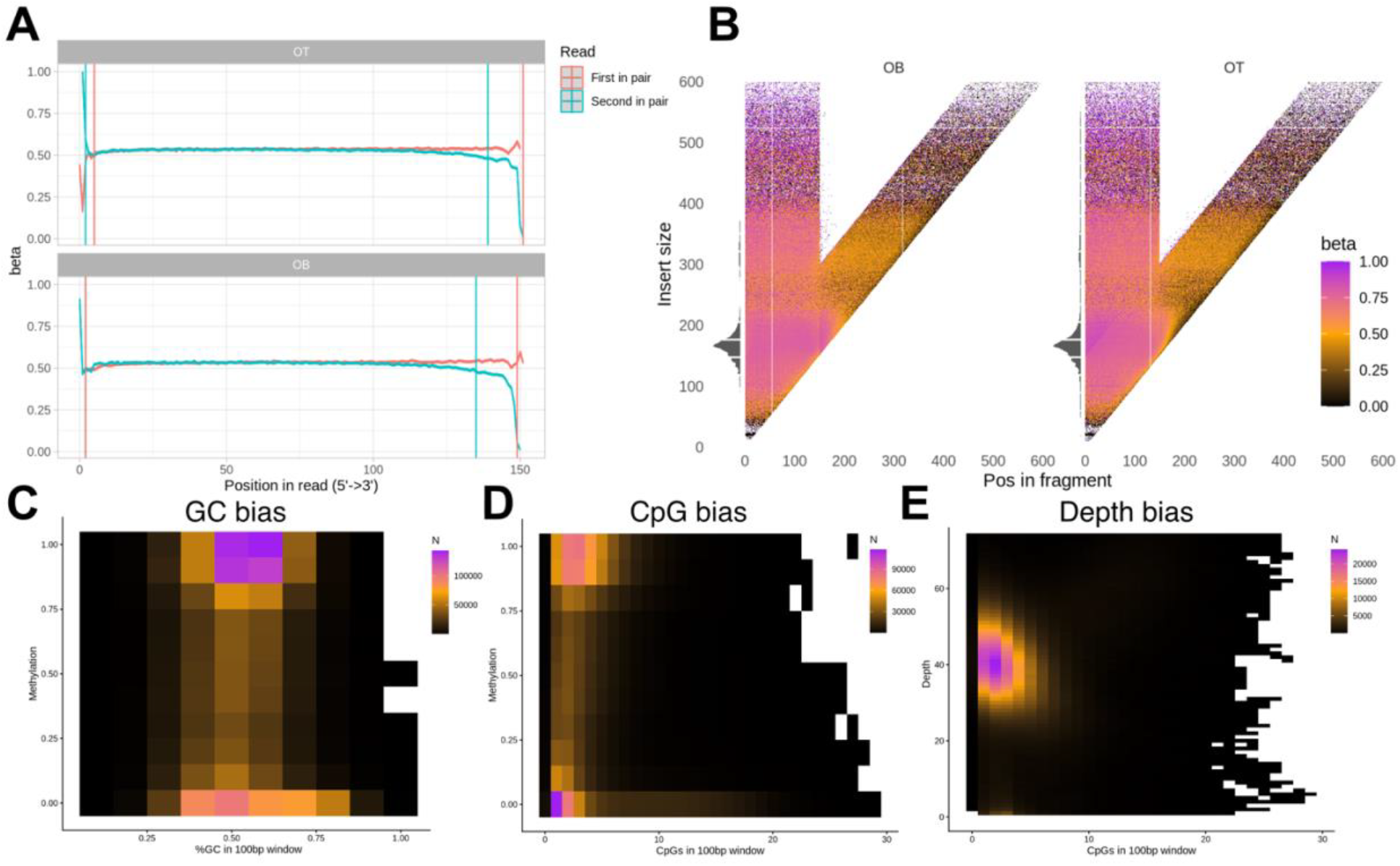
**(A)** M-bias plot for all reads on chr16 of the NA12878 TAPS library shown in Figure 2. **(B)** V-bias plot for a representative TAPS cfDNA sample of a cancer-free patient: we group fragments by their length and plot the average beta for CpGs at each fragment position. The right-hand side of each V reflects the second read in an Illumina read-pair, which has been reported to feature lower methylation. The histogram on the left side reflects insert-size distribution. **(C)** GC-bias: %GC in 100bp around the methylated position, **(D)** CpG bias: number of CpGs in 100bp around the methylated position, and **(E)** depth bias (effect of local CpG density on coverage) for the same NA12878 TAPS library as in Figure 2.

Finally, GC bias is a common phenomenon in most sequencing-by-synthesis approaches ^27^. The conversion of bases from C to T can change the GC content of reads and thus affect their propensity to amplify. We therefore also provide several plots to evaluate the effect of local GC content (Figure 5C) and the CpG density in a 100bp window on methylation (Figure 5D) and sequencing depth (Figure 5E).

## DISCUSSION

Rastair provides a robust framework to extract both genetic and epigenetic methylation information from mC→T data, such as those generated using Watchmaker’s TAPS+ or Illumina’s 5-Base chemistries. We have demonstrated that detection of genetic single-nucleotide variants at accuracy levels closely approaching standard whole-genome sequencing is indeed possible from such data, which sets mC→T methods apart from traditional bisulfite sequencing or methods such as EM-seq. Most experiments that aim to measure methylation with reasonable dynamic range per CpG will require sequencing depths in excess of 20x, at which point rastair’s accuracy in highly mappable parts of the genome is equivalent to that achieved by state-of-the-art algorithms on pure whole-genome sequencing data. This now enables the simultaneous measurement of epigenetic changes and genotype information. This could be leveraged to e.g. identify methylation-QTL relationships from a single assay, as we’ve demonstrated with the intriguing example of the rs1800734 SNP in the MLH1 locus.

CpG dinucleotides are the site of frequent C to T mutations and thus represent a hotspot of divergence within and between populations ^22^. This fact is frequently ignored when analysing methylation data, despite the fact that C to T transitions can mask as chemical conversion. Leveraging its fast and accurate genotyping model, rastair is able to detect such variant sites and correct the reported methylation levels to reflect the underlying genotype. This leads to a substantial shift (>10%) of estimated methylation levels (compared to existing tools that only report the number of C and T bases per position) at 68% out of the 543000 CpG sites across the genome that are affected by a genetic variant in NA12878. Similarly, few if any other methylation callers account for genetic variants that turn a non-CpG position into a CpG position. Such positions are often mis-interpreted as non-CpG methylation, since they show chemical conversion at a site where there is no CpG in the reference. We demonstrate that there are over 500,000 such positions in one individual, indicating that such “de-novo” CpGs represent a substantial amount of methylation information. Crucially, rastair provides methylation information at variant and de-novo CpG positions in two convenient and internally consistent formats: both the smaller and simpler bed output as well as the VCF output files contain information on all covered CpGs and the corresponding genotype-corrected methylation levels. Similarly, per-read output can be stored as a bed file that simplifies interactions with text-based downstream tools, or per-read methylation information can be stored directly inside the BAM alignment file. Care was taken to ensure standards compliance both for the VCF and BAM format to ensure that rastair can be easily integrated into existing downstream analysis pipelines.

With increasing output from new sequencing technologies such as Ultima Genomics or Roche’s Axelios system, the speed of tools to process these data is of growing importance. Rastair was carefully engineered with both performance and robustness in mind. It can process a whole-genome dataset sequenced to 45x in only 45 minutes using 32 CPU cores, faster than most other available tools. Where a compatible GPU is available, the runtime can be further reduced to less than 20 minutes, faster even than the hardware-accelerated DRAGEN tools which, according to Illumina’s run reports, take approximately 45 minutes to call variants on a similarly deeply sequenced file.

As TAPS+ and other mC→T methods gain popularity, we expect to continue to expand rastair’s feature set to match new use cases. We were careful to design the codebase in a modular fashion and to rigorously unit-test internal functions, in addition to automated integration tests using snapshots and bundled test data, to make it easier for external developers to contribute to rastair while ensuring ongoing correctness and avoiding regression errors.

Some areas of such future development are already defined on our roadmap. Currently, rastair does not attempt to determine local haplotypes for variants or methylation. Furthermore, rastair currently does not report insertion/deletion (indel) variants. This was a deliberate decision, given the intrinsic difficulty of calling indels accurately from short-read sequencing data: most variant calling algorithms have high concordance for SNPs but often low agreement on indel calls. Alignment artefacts are a major source of indel errors, so in a future iteration of rastair we plan to include a realignment step that accounts for methylation and subsequently tackle both phasing and indel calling on realigned reads. Finally, rastair currently does not deal with polyploid data and somatic variation, as encountered e.g. in cancer samples. This shortcoming will be addressed after indel and haplotyping have been implemented, so that haplotype structure can inform the somatic variant discovery.

The substantial simplification and reduction in cost brought about by new mC→T methods is driving a wave of new interest in methylation sequencing. These approaches are increasingly being used not just in an academic setting but also in a clinical and translational context, such as for cancer early detection. We hope that rastair, with its high performance, robust engineering and versatile outputs will provide the solid underpinning for this exciting future of epigenomics.

## CODE AND DATA AVAILABILITY

Source code for rastair is available from https://bitbucket.org/bsblabludwig/rastair. Rastair is also available as a conda package via bioconda, as a Docker image, and as a pre-built binary from https://www.rastair.com. Data are available upon request and will be uploaded to ENA before publication.

## ACKNOWLEDGEMENTS

This work was supported by the Ludwig Institute for Cancer Research. Computation used the Oxford Biomedical Research Computing (BMRC) facility, a joint development between the Centre for Human Genetics and the Big Data Institute supported by Health Data Research UK and the NIHR Oxford Biomedical Research Centre. The views expressed are those of the author(s) and not necessarily those of the NHS, the NIHR or the Department of Health. The authors would like to thank Watchmaker Genomics for generously providing data for training and testing.

## AUTHOR CONTRIBUTIONS

PH and BSB implemented rastair. LZ and ZE performed data analysis and contributed to the manuscript. BSB conceived the work, oversaw the analyses and wrote the manuscript.

