## Supplementary Information for "Rastair: an integrated variant and methylation caller"

### Supplementary Figures

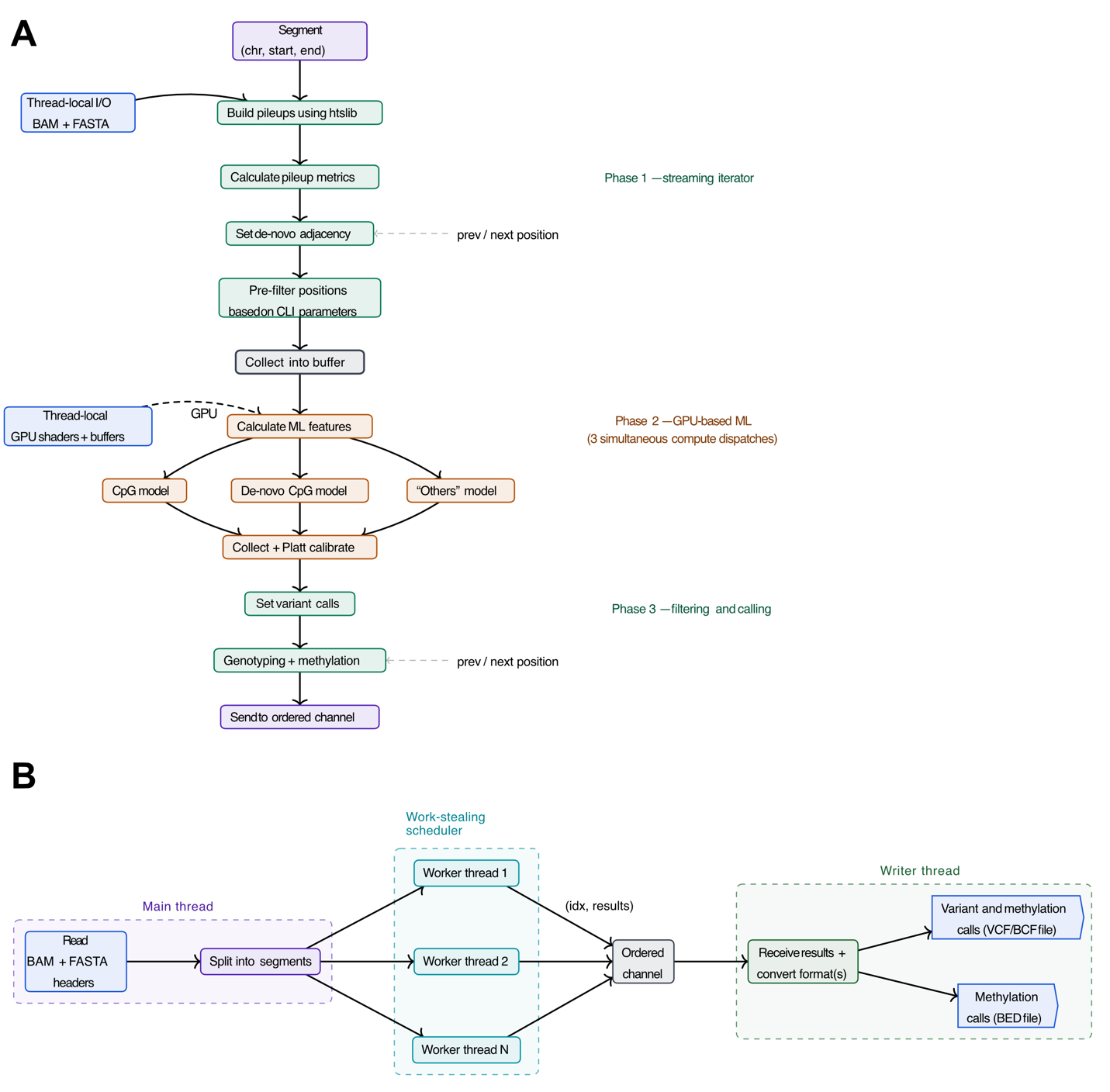

*Supplementary Figure 1 Outline of software architecture (A) Conceptual flow diagram of data processing for variants in each segment. (B) Flow diagram parallelisation across worker threads.*

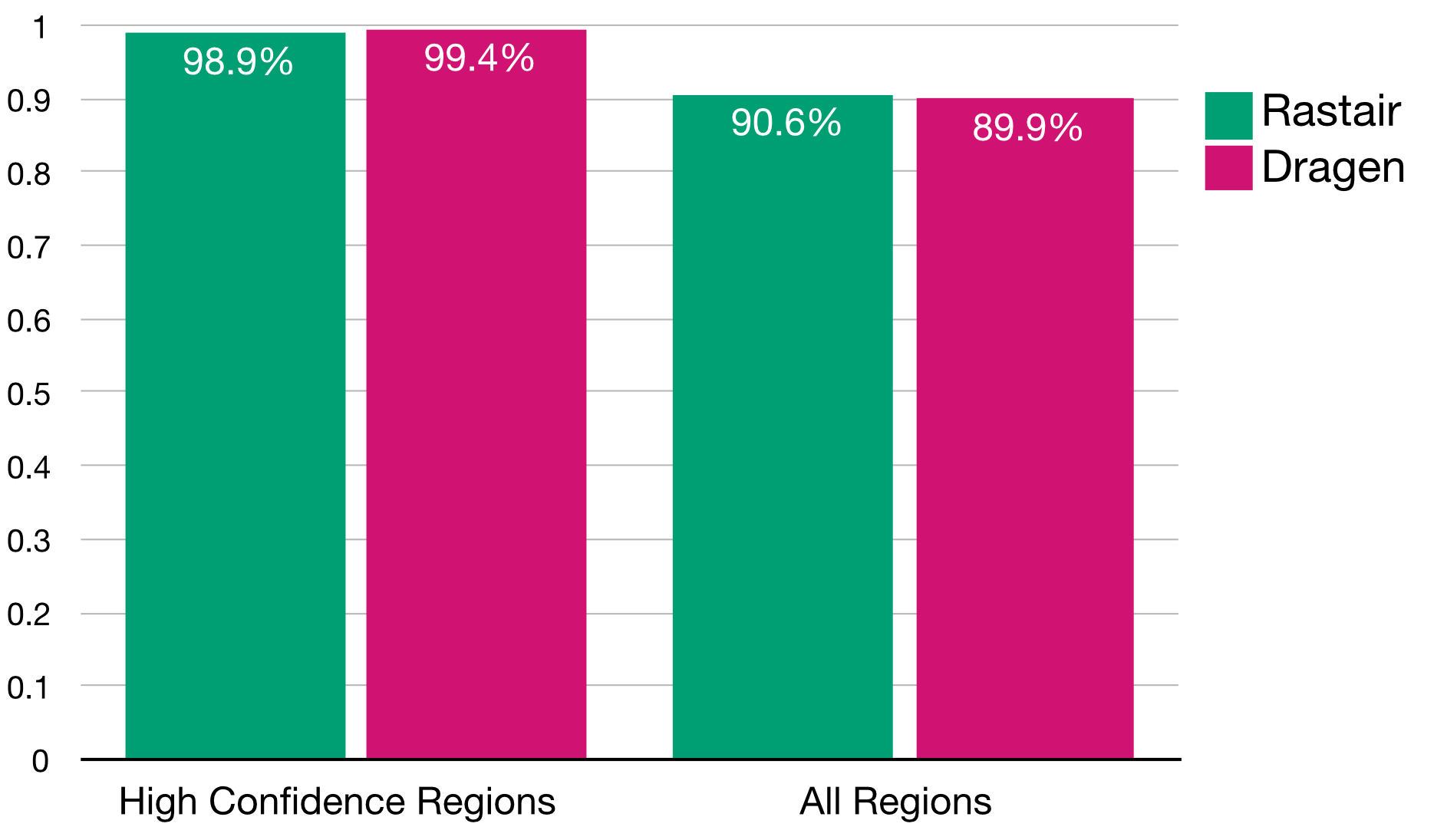

*Supplementary Figure 2 Comparison of DRAGEN and Rastair. Barplot showing F1 score of rastair (green) an DRAGEN (pink) in “High Confidence” regions (as defined by GIAB) vs all genomic regions.*

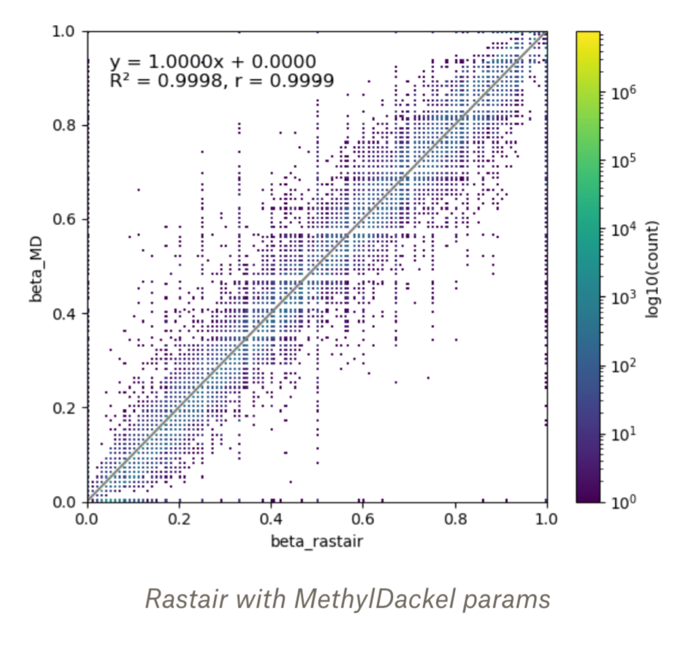

*Supplementary Figure 3 Matching filter settings of rastair and MethylDackel. Rastair was run with mapQ >= 10 and baseQ >= 5*

### Supplementary Tables

| **Caller** | **Tech** | **Coverage** | **CPUs** | **Memory** | **Mean (min)** | **SD (min)** | **Min (min)** | **Max (min)** | **Samples** |
| --- | --- | --- | --- | --- | --- | --- | --- | --- | --- |
| BISCUIT | TAPS | 10x | 32 | 256G | 12.4 | 0.1 | 12.3 | 12.6 | 3 |
| BISCUIT | TAPS | 30x | 32 | 256G | 14.4 | 0.2 | 14.2 | 14.7 | 3 |
| BISCUIT | TAPS | 45x | 32 | 256G | 15.3 | 0.2 | 15.1 | 15.5 | 3 |
| BISCUIT | TAPS | 5x | 32 | 256G | 10.6 | 0.4 | 10.2 | 11.0 | 3 |
| CLAIR3 | WGS | 10x | 32 | 64G | 280.4 | 4.2 | 276.2 | 284.6 | 3 |
| CLAIR3 | WGS | 30x | 32 | 64G | 187.3 | 7.4 | 182.7 | 195.8 | 3 |
| CLAIR3 | WGS | 45x | 32 | 64G | 226.7 | 4.4 | 223.8 | 231.7 | 3 |
| CLAIR3 | WGS | 5x | 32 | 64G | 176.5 | 3.3 | 174.4 | 180.4 | 3 |
| DEEPVARIANT | WGS | 10x | 32 | 64G | 77.6 | 1.6 | 76.5 | 79.5 | 3 |
| DEEPVARIANT | WGS | 30x | 32 | 64G | 98.2 | 2.3 | 95.6 | 99.7 | 3 |
| DEEPVARIANT | WGS | 45x | 32 | 64G | 114.9 | 1.7 | 113.6 | 116.8 | 3 |
| DEEPVARIANT | WGS | 5x | 32 | 64G | 67.4 | 1.5 | 65.9 | 68.8 | 3 |
| GATK | WGS | 10x | 32 | 64G | 414.3 | 1.7 | 412.5 | 415.8 | 3 |
| GATK | WGS | 30x | 32 | 64G | 726.4 | 6.6 | 720.1 | 733.2 | 3 |
| GATK | WGS | 45x | 32 | 64G | 898.6 | 19.5 | 880.6 | 919.4 | 3 |
| GATK | WGS | 5x | 32 | 64G | 301.0 | 13.3 | 287.6 | 314.1 | 3 |
| OCTOPUS | WGS | 10x | 32 | 128G | 100.1 | 2.9 | 96.8 | 102.0 | 3 |
| OCTOPUS | WGS | 30x | 32 | 128G | 181.3 | 5.1 | 175.4 | 185.0 | 3 |
| OCTOPUS | WGS | 45x | 32 | 128G | 233.6 | 2.5 | 231.1 | 236.2 | 3 |
| OCTOPUS | WGS | 5x | 32 | 128G | 69.3 | 1.2 | 68.6 | 70.7 | 3 |
| RASTAIR2 | TAPS | 10x | 32 | 96G | 15.7 | 0.1 | 15.6 | 15.8 | 3 |
| RASTAIR2 | TAPS | 30x | 32 | 96G | 32.7 | 0.7 | 31.9 | 33.2 | 3 |
| RASTAIR2 | TAPS | 45x | 32 | 96G | 44.2 | 0.6 | 43.6 | 44.7 | 3 |
| RASTAIR2 | TAPS | 5x | 32 | 96G | 10.5 | 0.1 | 10.4 | 10.6 | 3 |
| RASTAIR2_GPU | TAPS | 10x | 16 | 96G | 7.5 | 1.7 | 6.3 | 8.7 | 2 |
| RASTAIR2_GPU | TAPS | 30x | 16 | 96G | 14.7 | 2.2 | 13.3 | 17.2 | 3 |
| RASTAIR2_GPU | TAPS | 45x | 16 | 96G | 19.5 | 2.7 | 18.0 | 22.7 | 3 |
| RASTAIR2_GPU | TAPS | 5x | 16 | 96G | 5.5 | 0.4 | 5.0 | 5.8 | 3 |
| TVC | TAPS | 10x | 32 | 128G | 26.9 | 0.8 | 26.1 | 27.6 | 3 |
| TVC | TAPS | 30x | 32 | 128G | 80.8 | 3.5 | 76.9 | 83.6 | 3 |
| TVC | TAPS | 45x | 32 | 128G | 124.3 | 2.0 | 122.2 | 126.1 | 3 |
| TVC | TAPS | 5x | 32 | 128G | 13.4 | 0.6 | 12.7 | 13.8 | 3 |

### Supplementary Methods

#### Software architecture and performance design

Rastair is implemented as a highly-parallel command-line application written in Rust. It uses htslib for reading and writing most file formats (BAM, VCF, BCF), libdeflate from decompression, and the mimalloc allocator in combination with Rust's rich ecosystem to offer first-class performance, usability, and robustness.

To achieve its high runtime performance, Rastair splits the input data (or the user-select range) into segments whose processing a work-stealing scheduler distributes across a thread pool. The result of each processed segment is added to a ordered channel, i.e. out-of-order results will be buffered internally so that a single writer thread can receive the results sorted by their original positions and convert them seemlessly to the desired output format(s). Since segments are only given as a range of positions, each worker thread has its own thread-local readers for the input files (FASTA and BAM, both using indexes for random access). This combined with the buffered channel allows Rastair to saturate all available CPU cores with little to no lock contention and I/O bottlenecks.

Much of the calling accuracy is based on our usage of Random Forest models for variant evalulation. Profiling reveals this to be the most computation-intensive step in the pipeline of each worker with around 75% of time spend generating the model predictions (tested on a MacBook Pro M4 across multiple runs). Early in development we select the [biosphere](https://github.com/mlondschien/biosphere) library as the best fitting Random Forest library for Rastair. Rastair 2.1 is using a custom fork in which we added GPU-based inference using a flattened representation of the models running in compute shaders (using [wgpu](https://wgpu.rs/) to support Vulkan and Metal as well as DX12). This allows us to not only offload this step to dedicated hardware, but also to submit the inference calls for all three models (CpG, de-novo CpG, and other positions) simultaneously to the GPU. Similar to the thread-local readers, each Rastair worker thread has a thread-local GPU handle with models and buffers pre-allocated. Rastair also uses unified memory between CPU and GPU given hardware support.

We have continuously profiled the Rastair calling code using the [samply](https://github.com/mstange/samply/) profiler on macOS and Linux. This allowed us to find cases where we could change fundamental data structures to allow better cache line usage and stack-allocated structures. We also found performance bottlenecks in the rust-htslib wrapper library, which we fixed by adding read-level filtering and referenced views to reads.
